# Dynamics and Control of Phloem Loading of Indoleacetic Acid in Seedling Cotyledons of *Ricinus Communis* During Germination

**DOI:** 10.1101/042333

**Authors:** Imre A Tamas, Peter J Davies

**Author notes:** Deceased (Imre A Tamas). In commemoration of Ian Sussex, a noted scientist, colleague and friend whom I (PJD) overlapped at Yale University in 1966-1969.

## Abstract

During seed germination sugars and auxin are produced from stored precursors or conjugates respectively and transported to the seedling axis. To elucidate the mode of travel of IAA into the phloem a solution of [^3^H]indoleacetic acid (IAA), together with [^14^C]sucrose, was injected in the endosperm cavity harboring the cotyledons of germinating seedlings of *Ricinus communis*. Phloem exudate from the cut hypocotyl was collected and the radioactivity recorded. Sucrose loading in the phloem was inhibited at higher IAA levels, and the rate of filling of the transient pool(s) was reduced by IAA. IAA was detected within 10 minutes, with the concentration increasing over 30 min and reaching a steady-state by 60 min. The kinetics indicated that phloem loading of IAA involving both an active, carrier-based, and a passive, diffusion-based component, with IAA traveling along a pathway containing an intermediary pool, possibly the protoplasts of mesophyll cells. Phloem loading of IAA was altered by sucrose, K^+^ and a range of non-specific and IAA-specific analogues and inhibitors in a manner that showed that IAA moves into the phloem from the extra cotyledonary solution by multiple pathways, with a carrier mediated pathway playing a principal role.

**Summary Statement:** Indoleacetic acid is transported from the peri-cotyledonary space into the phloem of germinating *Ricinus* seedlings by both trans-membrane carriers and diffusive pathways, with the cells of the cotyledons forming an intermediate reservoir.

## Introduction

Seeds contain not only food reserves in the form of oils or starch but also hormones on a precursor or conjugate form (Normanly et al., 2010). In the case of the auxin (indole-3-acetic acid [IAA]) these can be on the form of indoleacetyl aspartate, IAA inositol or glycoside conjugates, or complexes thereof, or peptide conjugates. During germination the oil or starch reserves and converted into sucrose for transport to the developing embryo, whereas IAA is released from conjugates (Morris et al., 2010). The mechanism by which IAA enters the transport stream is currently unknown. Germinating seeds of castor bean (*Ricinus communis*) form an ideal model system to address this question because the reserves are held in the endosperm and during germination the solubilized reserves are taken up by leaf-like cotyledons located in the center of the endosperm tissues. These cotyledons function initially as absorptive organs, and only later emerge to function as the first leaves of the seedling. As these cotyledons have no cellular connection to the endosperm reserves, both reserve materials and hormones alike must be solubilized, transported into the peri-cotyledonary space, and then taken up by the cotyledons prior to transport to the rest of the seedling. Cotyledons of the Ricinus seedling readily take up solutes from the incubation medium via the whole blade surface (Komor et al., 1991). When the hypocotyl is severed the seedlings exude phloem sap from the cut hypocotyl (ibid.). The imbibed, germinating seedlings of caster bean are large enough for the easy application of tracer materials and pharmaceutical modifying agents into this peri-cotyledonary space, so that germinating Ricinus seedlings make an excellent system to address these questions. In addition Ricinus seedlings have been extensively utilized for studies on sucrose uptake by the cotyledons, so the characteristics of sucrose uptake are well known (Komor et al., 1991; Orlich et al., 1998; Orlich and Komor, 1992)

### Sucrose Uptake by the Phloem in Ricinus Seedlings

During germination and the first few days of growth sucrose, derived from stored materials in the endosperm, is absorbed by the cotyledons from the surrounding medium and thence transported into the phloem (Weig and Komor, 1996). Sucrose is the major solute in the phloem (Komor et al., 1991) and there is an accumulation of sucrose in the phloem to a concentration of 270 mM, exceeding that in the surrounding medium, indicative of active transport (Kallarackal et al., 1989). At least 50 % of sucrose is loaded by the apoplastic pathway without involvement of the symplastic route (Komor et al., 1991). The pH dependence of sucrose uptake (with an optimum of pH 5) (Weig and Komor, 1996), combined with the alkalization of the apoplast and membrane depolarization of mesophyll cells is indicative of H^+^/sucrose cotransport (ibid.; (Koehler et al., 1991) There is also a direct uptake of sucrose by the sieve tube-companion cell complex from the apoplast (ibid.). About half of the sucrose exported is loaded into the phloem directly from the apoplasm, while the other half takes the route via the mesophyll. Mesophyll-derived sucrose is released into the apoplasm adjacent to the phloem prior to loading into the phloem (Orlich and Komor, 1992), so that loading into the phloem is by both a direct and an indirect apoplasmic pathway as well as symplasmic loading

Williams et al. (1990), using cotyledon-derived plasma-membrane vesicles, provided strong evidence for a sucrose-proton symporter system in the plasma membrane of cells of Ricinus cotyledons. Sucrose uptake had a Km of 0.87mM (Komor et al., 1991) was stimulated by a pH gradient with a pH optimum of pH 6.5, was inhibited by vanadate, the sulfhydryl reagent p-chloromercuribenzenesulfonate (PCMBS) and the protonophore CCCP, and showed strong specificity for ATP as a substrate (Williams et al., 1990). A sucrose-carrier gene was found to be expressed in the cotyledons of Ricinus seedlings at a similar level at germination and 3-6 days after germination, with the greatest expression in the lower epidermal layer and the phloem, consistent with an active loading role for these cells (Bick et al., 1998b).

The energy source for the process of phloem loading in those plants utilizing sucrose-proton cotransport is an electrochemical potential gradient maintained through the active extrusion of protons by H^+^ pumps into the apoplastic free space (Hutchings, 1978a). In excised cotyledons of *Ricinus* seedlings, externally applied sucrose readily enters the apoplast whence it is actively loaded into the sieve element/companion cell-complex by an H^+^/sucrose cotransport system (Hutchings, 1978a). A significant portion of applied sucrose may however be taken up by the mesophyll and passed on to the phloem via symplastic flow (Orlich et al., 1998). Sucrose uptake by excised *Ricinus* or soybean cotyledons shows a biphasic response to sucrose concentration. At low external levels, sucrose uptake operates as a high affinity, carrier based process, characterized by low rates of H^+^ and sucrose influx. With increasing sucrose concentration a linear, diffusion-like phase becomes predominant between 20 and 50 mM sucrose, showing a diminished dependence on net H^+^ influx and a consequent sharp decline in the stoichiometry of H^+^: sucrose (Delrot and Bonnemain, 1981; Hutchings, 1978a; Komor, 1977; Kriedemann and Beevers, 1967; Lichtner and Spanswick, 1981).

When potassium ions are included along with sucrose in the incubation medium, there is a bimodal effect on sucrose loading depending on K^+^ concentration: stimulation of loading at the lower range (generally below 10 mM K^+^), and inhibition at higher levels (Hutchings, 1978b; Komor et al., 1977; Schobert et al., 1998; Van Bel and Koops, 1985). At the lower range, K^+^ influx will allow a modest rate of discharge of the pH-gradient, which results in a faster recirculation of H^+^ and enhanced sucrose loading without a major drop in membrane potential. At increasingly higher levels, K^+^ influx will depolarize the plasma membrane causing a concentration-dependent decrease in transport activity.

Amino acids are also taken up by proton mediated carriers. Glutamine was taken up by plasma membrane vesicles with a Km of 0.35mM and similar sensitivity as sucrose to both PCMBS and CCCP (Williams et al., 1990). Bick et al. (1998a) found genes for two putative amino acid carriers to be abundantly expressed in the cotyledons.

### Long Distance Auxin Transport

Auxin is naturally exported from source leaves in the phloem (Baker, 2000a; Morris et al., 2010). Ricinus phloem sap, collected via incisions into the inner bark, has been found to contain 13 ng ml^−1^ of IAA in phloem sap (as analyzed by gas chromatography-mass spectrometry) (Baker, 2000b) and this sap provides one of two sources of auxin to the rest of the plant, the other being cell to cell polar auxin transport; the xylem had only a small fraction of this amount (ibid.). IAA applied to mature pea leaflets was initially exported via the phloem as detected by aphids feeding on the stem or recovery in exudates collected from severed petioles (Cambridge and Morris, 1996), and endogenously-produced IAA was found in the phloem exudate from excised pea leaflets at a production rate of 7.7 pg leaflet^−1^ h^−1^(Jager et al., 2007), though as the volume was not recorded the concentration cannot be calculated. Mature leaves are therefore one source of the IAA in the basipetal transport stream. After a period of hours applied IAA exported from leaves in the phloem was found transferred into the extravascular polar auxin transport pathway though reciprocal transfer from the polar auxin transport stream into the phloem probably does not occur (Cambridge and Morris, 1996).

Polar auxin transport relies in a pH gradient-driven weak-acid passive uptake and carrier mediated uptake, combined with specific IAA transporters located in the base of each cell (Morris et al., 2010), leading to an iteration from cell to cell and thus transport in a basipetal direction. IAA loaded into the phloem will be subject to the factors that influence phloem translocation (ibid.). A principal such factor is loading into the phloem. Several pathways of phloem loading exist, namely simple concentration-mediated diffusion, transmembrane proton co-transport, and a polymer trap mechanism, which may operate singly or in combination, and for sugars this varies with the species (Rennie and Turgeon, 2009). The mechanism of IAA entry to the phloem is unknown. Previous work on IAA uptake into stem segments has revealed various mechanisms of entry into the transport stream, and the way that these can be distinguished pharmacologically (Davies and Rubery, 1978). The aim of this work was to ascertain the mechanism and extent of IAA transport from the seed source into the phloem of *Ricinus* seedlings. *Ricinus* seedling cotyledons represent a natural way of investigating this phenomenon because there is a lack of cellular connection between the endosperm and the cotyledons (Komor et al., 1991) such that the application of sucrose and IAA mimics the natural pathway that already exists. This pathway into the cotyledonary cells and on into the phloem can be elucidated for IAA using techniques similar to those already used to examine sucrose movement for extra-cotyledonary sucrose into the phloem of the *Ricinus* seedlings.

## Results and Discussion

### Simultaneous transport of differentially labeled IAA and sucrose in phloem: Effect of IAA concentration

After a buffered medium containing both [^3^H]IAA and [^14^C]sucrose was injected in the endosperm cavity harboring the cotyledons, both substances were recovered in the phloem exudate from the severed end of the hypocotyl stump about 1 cm below the cotyledons and endosperm (Fig. 1). A broad range of IAA concentrations from 0.0016 to 20 mM was tested, all with identical [^3^H]IAA content. Freely exuding phloem sap was collected at 10 min intervals up to 1 h. The volume exuded in 10 min ranged from 2 to 6 μL. The volume and amount of radioactivity recovered were highly variable, from seedling to seedling, though largely consistent over an hour within any one seedling. According to Komor and co-workers (Komor et al., 1991) the flow rate is determined by the rate of phloem loading, the osmotic water uptake, the resistance of the sieve tubes and the percentage of open phloem; factors that account for variability of the exudation rate include the seedling age and stage of development, handling and quality of cut, and the composition of the medium. Despite differences in the exudation rate, the solute composition and concentration of the exudate is reported to be very similar between individual seedlings (ibid.). When our results were corrected for the volume of solution recovered, i.e., as the concentration (in %) relative to the injected concentration, patterns became clear. At all applied levels IAA was detected within the first 10 minutes, and the relative concentration of IAA appearing in the exudate increased at similar rates for the first 30 minutes (Fig. 2A). Over the next 30 minutes the IAA exudation rate gradually came to an apparent steady state level. This pattern may be obtained when the applied radiolabeled substance passes through one or more transient pools of defined size in the transport pathway to the site of phloem loading. When the substance entering the phloem has reached a steady fraction of the concentration in the injection medium, the recovered concentration also attains a steady state. The fact that the relative steady state levels for different IAA applied concentrations were in close proximity indicates that IAA loading in the phloem has stabilized at very different IAA levels - stretching over four orders of magnitude - roughly in proportion to the applied concentration, except it was somewhat less than that at the highest concentration (20 mM). These observations show that IAA transport in phloem can occur efficiently over a very wide concentration range, reflecting perhaps the activity of a complex transport system with multiphasic kinetics (Komor et al., 1977). More specifically, the results suggests the operation of a diffusive, ‘linear’ component at the higher levels (Komor et al., 1977; Lichtner and Spanswick, 1981), analogous perhaps to the mode of sucrose transport at varying sucrose concentrations (Fig. 6B). The fact that the relative IAA transport rate at the highest level of applied IAA (20 mM) was somewhat less than linear (Fig. 2A) suggests that IAA concentration is not the sole factor controlling the loading process.

**Fig. 1.**
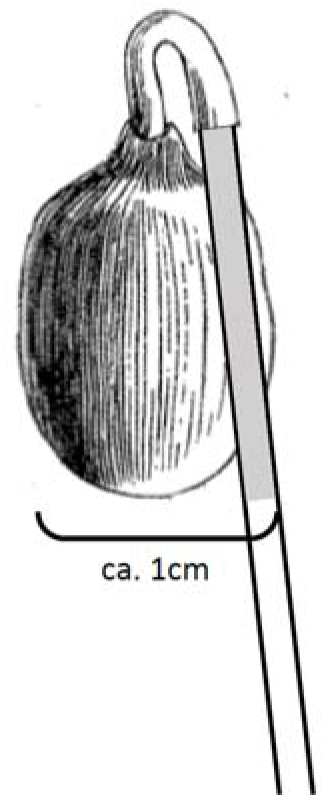
Diagram of the treatment method. The hypocotyl of hydrated 6-day old Ricinus seedlings was cut off leaving a 1cm stump, which was placed against a calibrated 10μL capillary tube. Five μL of injection medium containing the radiolabeled substances to be transported was injected into the endosperm cavity between the cotyledons. The capillary tube with exuded phloem sap was replaced every 10 min, the volume recorded, and the contents counted for radioactivity.

**Fig. 2.**
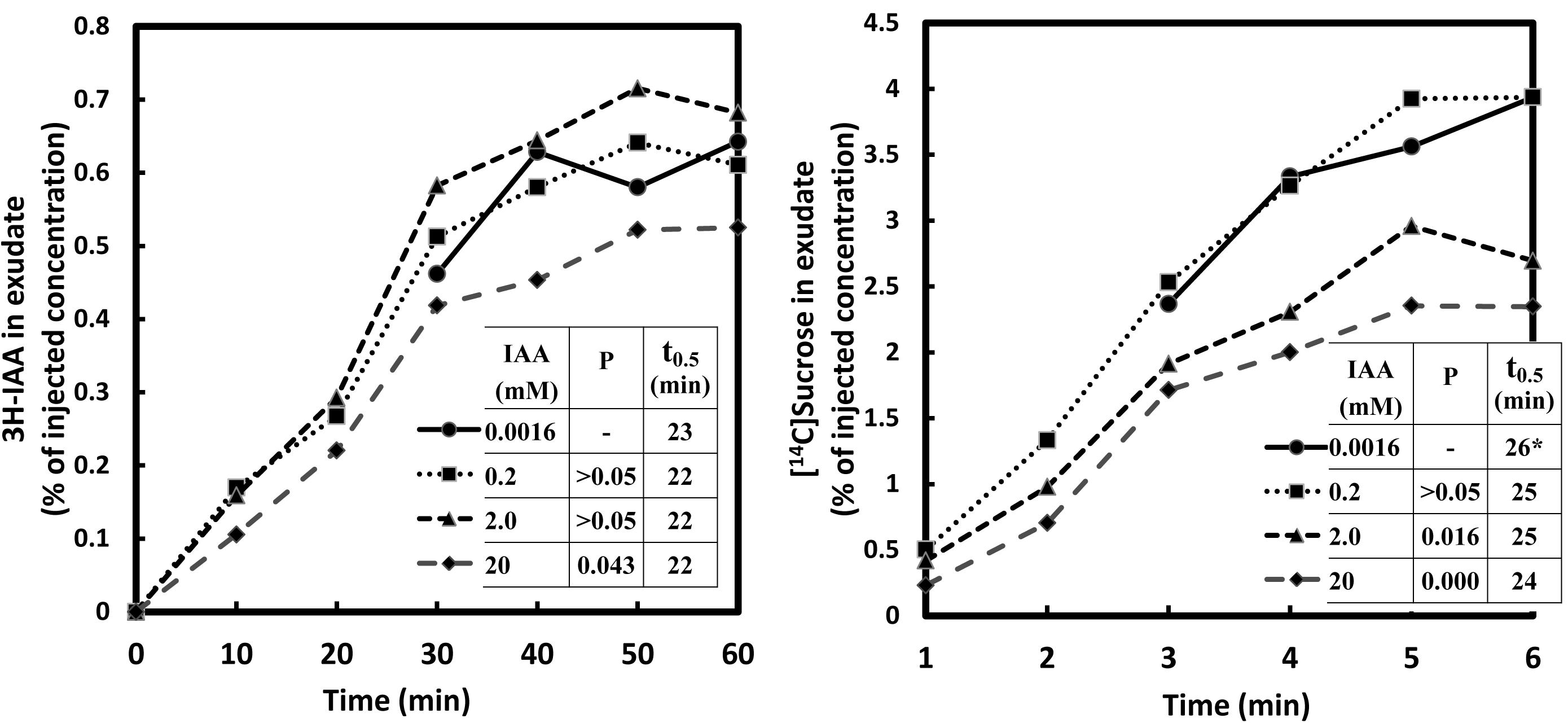
Simultaneous transport of [^3^H]IAA and [^14^C]sucrose from the cotyledons to the hypocotyls of germinating seedlings of *Ricinus communis* at various levels of IAA. Each of four injection media contained 30 mM MES buffer at pH 6.3, 20 mM Na^+^, 1.48 MBq mL^−1^ [^3^H]IAA, and 0.74 MBq mL^−1^[^14^C]sucrose. Non-radioactive IAA was added to adjust the concentration levels to 0.0016, 0.2, 2.0, and 20.0 mM. The concentration of injected sucrose was 0.35 mM. Transported IAA (A) or sucrose (B) concentration found in the phloem exudate (from the collected radioactivity per unit volume of exudate) is expressed as the percentage of the applied IAA or sucrose concentration respectively. The data represent each individual 10 min collection and are not cumulative. The variable *t0.5* is the time (in minutes) at ½-steady-state (or at ½-maximum) concentration attained within 1 h of transport. If no steady-state is apparent, the *t0.5* value is marked with an asterisk. The data are the combined results of three independent experiments, and represent the means of at least ten replicate measurements.

The [^14^C]sucrose was injected into the endosperm cavity at a sufficiently low level (0.35 mM) that it would have a negligible effect on the native sucrose concentration, estimated to be about 90 mM (Kriedemann and Beevers, 1967). Transported [^14^C]sucrose concentration in the phloem exudate at 1 h, expressed as the percentage of the injected sucrose concentration, was about 4% at the two lower applied IAA levels (Fig. 2B). The corresponding transported sucrose concentration was only about 2.5% in the presence of 2.0 or 20.0 mM IAA, indicating that sucrose loading in the phloem is slightly subject to inhibition at higher, non-physiological IAA levels. As the time-dependent changes in transported sucrose concentration revealed, the trend was at or near linearity for the lowest applied IAA level throughout the one hour run (the *t0.5* value [the time (min) required for the transported substance to attain the ½-steady-state level in the phloem exudate] is marked with an asterisk to note that a steady-state has not been attained). However, the trend became progressively more sigmoid with increasing IAA levels: at 20 mM IAA there was in the first 20 min a much slower appearance of the labeled sucrose in the exudate indicating that the rate of filling of the transient pool(s) was reduced by IAA. Perhaps as a consequence, sucrose transport at 20 mM applied IAA came to a steady-state at a concentration much below that of the lowest IAA level (Fig. 2B).

### NAA competes with IAA transport in the phloem

The synthetic auxin α-naphthaleneacetic acid (NAA) is analogous to IAA in many of its physiological properties, including the ability to serve as a competitive substrate for the auxin efflux carrier. However, in contrast to IAA, NAA has only marginal affinity for the auxin influx carrier (Delbarre et al., 1996; Marchant et al., 1999). Externally applied NAA enters cells by diffusion. On the basis of these attributes, we selected NAA as a diagnostic probe to test whether IAA loading into the phloem requires auxin efflux carrier activity. We measured the transport of [^3^H]IAA (0.78 μM) injected in combination with NAA at concentrations of 0.0, 0.1, 1.0, and 10.0 mM. Analysis of the collected samples of phloem exudate showed that [^3^H]IAA transport was inhibited in plants treated by NAA, the effect being most highly expressed at both 0.1 and 10.0 mM (Fig. 3A). In the affected plants progress toward a steady-state was slower, and it was reached at a lower IAA concentration. The competitive effect of NAA suggests that in *Ricinus* cotyledons the passage of [^3^H]IAA through the loading pathway is facilitated by auxin efflux carriers. Basipetal transport of IAA, involving efflux carriers, may take place in files of parenchyma cells that are closely associated with minor veins in developing leaves as described in the next section (Aloni, 2010; Mattsson *et al.,* 1999; Mattsson *et al.,* 2003).

**Fig. 3.**
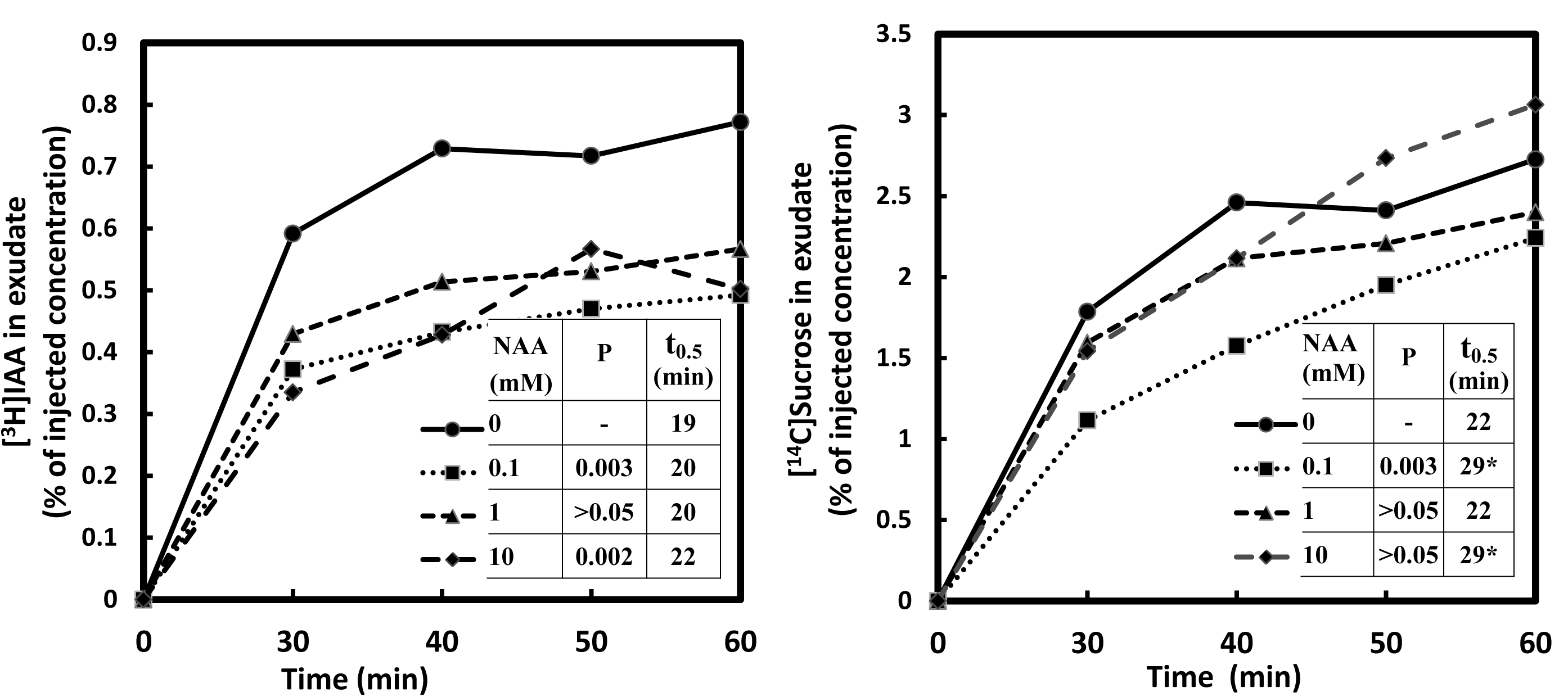
Transport of [^3^H]IAA (A) and [^14^C]sucrose (B) in the presence of NAA. The injection media contained 0.74 MBq mL^−1^ [^3^H]IAA (0.78 μM) and 0.19 MBq mL^−1^[^14^C]sucrose (8.3 μM). NAA concentrations were 0.0, 0.1, 1.0, and 10.0 mM. Other conditions or comments were as in Fig. 2.

To test whether sucrose transport may also be altered in the presence of NAA, [^14^C]sucrose at 8.3 μM was included in the injection medium along with [^3^H]IAA. The effect, compared to that on IAA transport, was much less clear. There was some reduction in sucrose transport but only at the lowest (0.1 mM) level of NAA (Fig. 3B). This is an interesting result as it seems to contradict the inhibiting effect of IAA on sucrose transport (Fig. 2B).

### Phloem loading of IAA is stimulated by the auxin transport inhibitor triiodobenzoic acid

Phloem transport of [^3^H]IAA from cotyledons of *Ricinus* seedlings was stimulated at both 20 and 100 μM 2,3,5-triiodobenzoic acid (TIBA) (Fig. 4A). The results suggest that a TIBA-enhanced IAA accumulation in auxin-transporting tissues caused a diversion of IAA flow toward the phloem, or an inhibition of lateral efflux from the final sieve tubes. This conclusion is supported by published evidence indicating that TIBA and other auxin transport inhibitors cause auxin accumulation in cells (Davies and Rubery, 1978); that auxin accumulation results in lateral transport between neighboring cells or tissues (Nicolas et al., 2004); that lateral auxin transport may be an integral component in auxin signaling pathways; and that the direction and rate of lateral transport is determined by the prevailing auxin concentration gradient within the transport pathway (ibid.). The role of TIBA as an inhibitor of the auxin efflux carrier *PIN1* has been extensively documented in studies on polar auxin transport (Morris et al., 2010). Applied TIBA inhibits the basipetal release of auxin by cells in the polar transport pathway, thereby causing auxin accumulation (Davies and Rubery, 1978). With rising auxin concentration, the lateral release of auxin to neighboring tissues is enhanced thus altering the relative flux among different transport pathways. Evidence for such a mechanism is contained in a study on vascular patterning in *Arabidopsis* leaves showing that auxin transport provides the controlling signal for both the initiation and the subsequent development of vascular strands in growing leaves. Basipetal auxin transport originating in the tip of young leaf primordia will set the location of the primary vein by inducing the formation of a line of procambial cells. Continuing basipetal auxin transport in the fascicular cambium of the developing vein, together with lateral auxin flow from neighboring cells, controls the ultimate size and composition of the vein (Aloni, 2010; Mattsson *et al.,* 1999; Mattsson *et al.,* 2003) (such a cambium would be restricted to the major veins as the minor veins are too small consisting only of a very few cells, though parenchyma cell(s) may be included in these smaller veins.) Cotyledons of *Ricinus* seedlings also possess an extended bundle sheath that serves as a transport tissue and a temporal sink for assimilates (Rutten et al., 2003) and possibly also auxin. The rate of lateral auxin flow varies with the concentration gradient, which is maintained by the drainage capacity, or “sink effect”, of the vein.

**Fig. 4.**
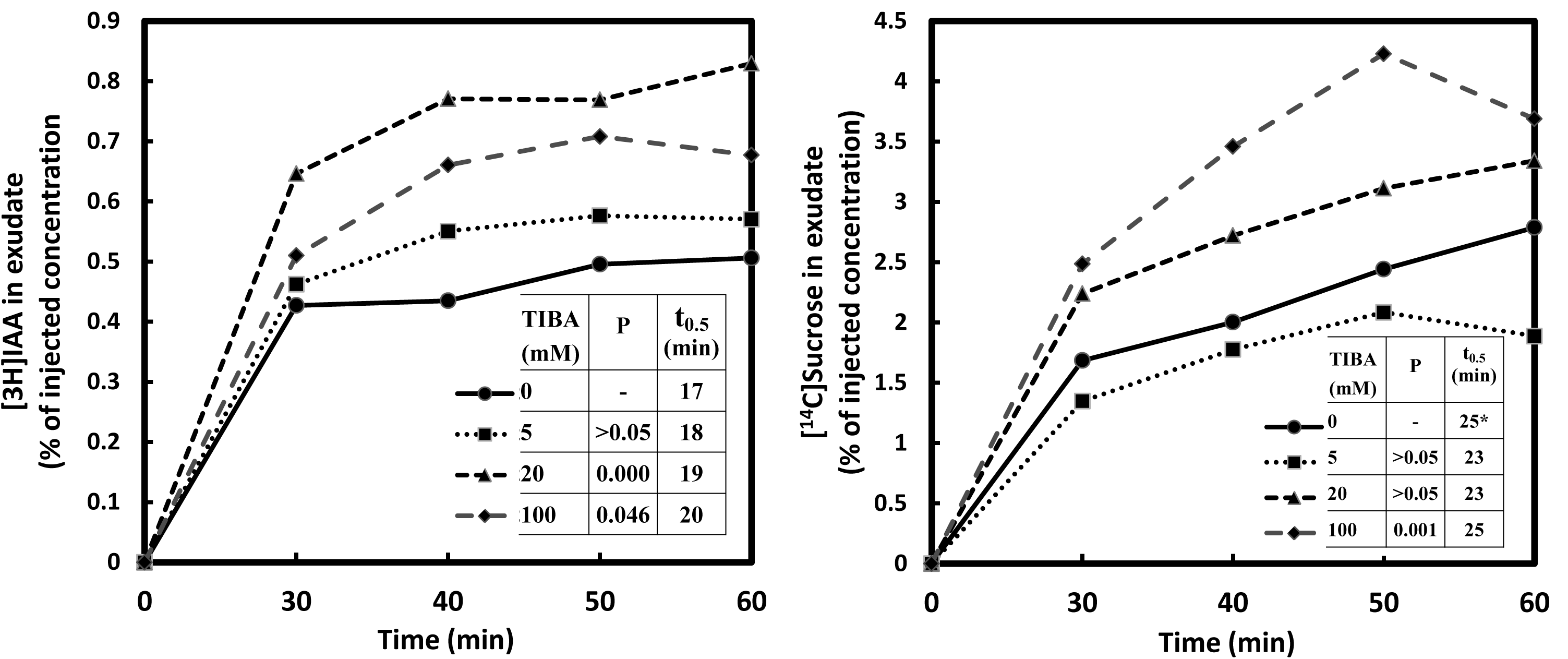
Effect of TIBA on the transport of [^3^H]IAA (A) and [^14^C]sucrose (B). The injection media contained 0.74 MBq mL^−1^ [^3^H]IAA (0.82 μM) and 0.19 MBq mL^−1^[^14^C]sucrose (7.44 μM). TIBA concentrations were 0, 5, 20, and 100 μM. Other conditions or comments were as in Fig. 2.

Several lines of evidence support the notion that auxin transport inhibitors can alter the rate of lateral auxin efflux from cells. Results by Mattsson *et al*., (1999) suggest that in developing *Arabidopsis* leaves the lateral movement of auxin toward the vascular strands was enhanced significantly by treatment with NPA or TIBA as shown by the increased width of the developing veins. In transgenic *Arabidopsis* seedlings, subjected to gravity or light stimulation, there was a tropic bending response of the hypocotyl which occurred concurrently with an elevated expression of the synthetic *DR5::GUS* auxin reporter gene on the more elongated side of the hypocotyl. The effect was attributed to the lateral relocation of the auxin efflux regulator gene *PIN3* as shown by immunogold electron microscopy (Friml et al., 2002). Plants receiving gravity or light stimulation in the presence of NPA failed to show asymmetric *DR5::GUS* expression or tropic curvature. NPA prevented the actin-dependent lateral redirection of auxin by inhibiting the relocalization of the *PIN3* protein in the cell plasma membrane.

Much of our knowledge about TIBA relates to its role in inhibiting polar auxin transport. There is, however, accumulating evidence that its action is more broadly based through a general influence on cellular protein trafficking (Geldner et al., 2001). TIBA and other auxin transport inhibitors were shown to retard auxin transport by blocking *PIN1* cycling, and also to interfere with the trafficking of plasma membrane H^+^-ATPase and of other proteins. In the present work we show that in plants treated with 100 μM TIBA [^14^C]sucrose transport is enhanced, as is [^3^H]IAA transport (Fig. 4B). Conceivably, the localization of sucrose transporters could also be altered by TIBA as these proteins are degraded and turned over.

### Effect of potassium ion and sucrose concentration

The uptake and phloem loading of sucrose is known to be controlled by a diverse set of internal or externally applied factors including inorganic ions, pH, substrate concentration, as well as reagents for probing metabolic or transport activity (Komor, 1977; Maynard and Lucas, 1982; Schobert et al., 1998; Williams et al., 1992). Given the complex role that sucrose and potassium ions seem to play in phloem function, we examined the effect of these factors on IAA transport. Phloem input and transport rates of IAA and of sucrose were measured together at varying sucrose concentrations, with or without 20 mM K^+^ present in the injection medium; in the latter case 20 mM Na^+^ was substituted for K^+^.

With the inclusion of 20 mM K^+^ in the injection medium the pattern of sucrose transport was altered compared to that without K^+^. With 0.02 mM sucrose, the sucrose content of the exudate was about 0.9% at the end of the 1 h run (Fig. 6B), a value less than half of that obtained without K^+^ (Fig. 5B). Therefore, 20 mM K^+^ in the medium was inhibitory for sucrose transport, a finding in agreement with published results (Van Bel and Koops, 1985). Also at this low applied sucrose level, the presence of potassium caused a shift in the time course from a largely linear to a strongly sigmoid shape, perhaps indicating a shift toward a longer loading pathway. With potassium present there was no significant difference in the relative sucrose transport rates at the three applied sucrose levels, so that the sucrose flux increased in proportion to the applied concentration, suggesting that transport activity at the two higher levels was predominantly in its linear, non-saturable phase (Fig. 6B). Also, with higher applied sucrose levels the value of *t0.5* was much increased, indicating a lengthening loading pathway and a strong upward trend in the transient pool size (Fig. 6B); this may mean that a relatively greater portion of transported sucrose was passing through the mesophyll on its way to the phloem.

**Fig. 5.**
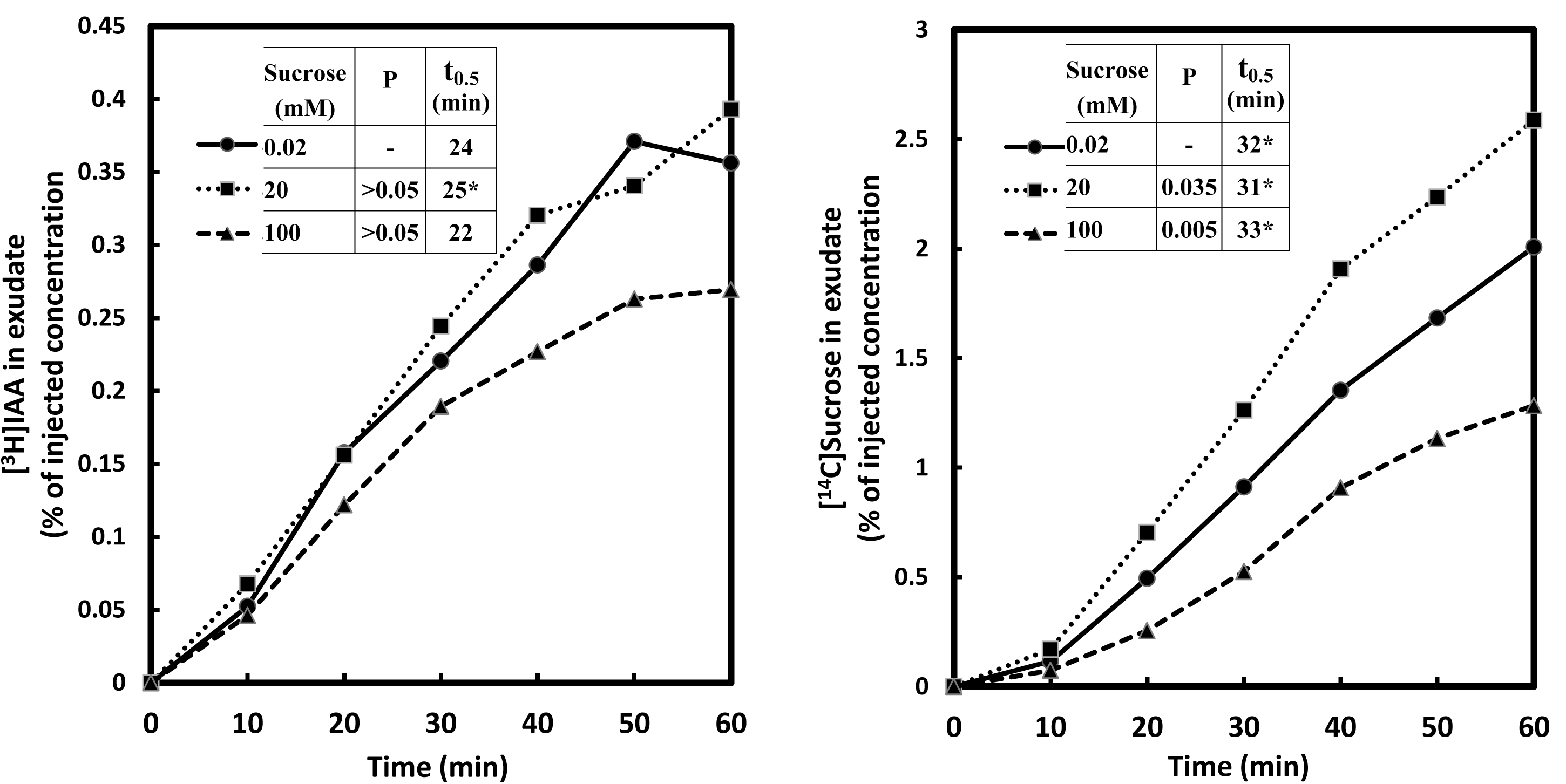
Transport of [^3^H]IAA (A) and [^14^C]sucrose (B) at various levels of sucrose. Each of the injection media contained 30 mM MES buffer at pH 6.3, 20 mM Na^+^, 0.71 MBq mL^−1^ [^3^H]IAA (0.76 μM), and 0.63 MBq mL^−1^[^14^C]sucrose. Non-radioactive sucrose was added to adjust concentration levels to 0.02, 20, and 100 mM. The data are combined from two experiments, and the means are from eight replicate measurements. For other conditions or comments see Fig. 2.

**Fig. 6.**
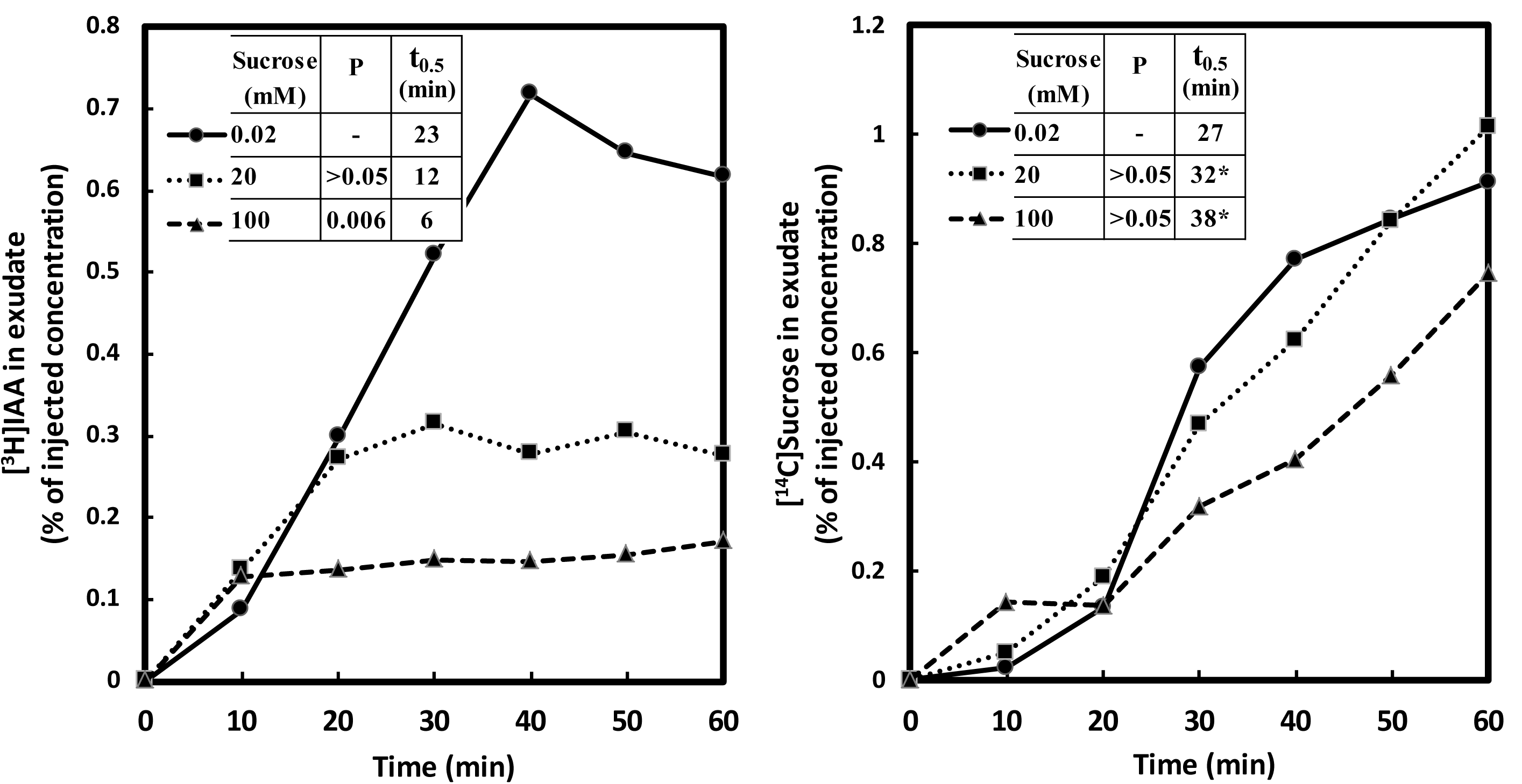
Transport of [^3^H]IAA (A) and [^14^C]sucrose (B) at various levels of sucrose in the presence of potassium ion. Each of the injection media contained 30 mM MES buffer at pH 6.3, 20 mM K^+^, 0.91 MBq mL^−1^ [^3^H]IAA (0.98 μM) and 0.524 MBq mL^−1^[^14^C]sucrose. Non-radioactive sucrose was added to adjust concentration levels to 0.02, 20, and 100 mM. For other conditions or comments see Fig. 5.

In the absence of potassium ions, the effect of sucrose concentration on transport rates was either insignificant, as in the case of IAA (Fig. 5A), or inconsistent, as in the case of sucrose (Fig. 5B). The inclusion of 20 mM K^+^ in the injection medium evoked a set of correlated changes in IAA transport (Fig. 6A) that provide a striking contrast to the results obtained in the absence of K^+^ (Fig. 5A). At the lowest sucrose level, the amount of IAA nearly doubled after 1 h of transport due to the presence of potassium, presumably resulting from an enhancement of the plasma membrane H^+^-gradient with K^+^ acting as a counter-ion. Whereas the steady-state concentration of transported IAA in the phloem exudate in the presence of K^+^ was about 0.7 % at 0.02 mM sucrose, it was reduced to about 0.3 % and 0.15 % at 20 mM and 100 mM sucrose respectively (Fig 6A). In addition the t0.5 values in the presence of K^+^ declined from 23 min at 0.02 mM sucrose to 12 and 6 min at 20 mM and 100 mM sucrose respectively (Fig 6A). These responses are in agreement with, and are explained by the combined effects of sucrose and K^+^ on phloem loading previously described. Therefore, the following conclusions may be drawn from the interactions of K+ and sucrose on IAA loading into the phloem (Fig. 5A and 6A): 1) The stimulation of IAA loading by K^+^ suggests that the IAA carrier was in its high affinity phase at the applied concentrations of 20 mM K^+^ and 0.02 mM sucrose, and therefore the load-enhancing range of K^+^ for the IAA carrier must be wide enough to include the 20 mM level; 2) The degree of sensitivity of IAA loading to the depolarization of the plasma membrane is correlated with sucrose concentration; 3) A t0.5 value may be taken as a semi-quantitative measure of the collective size of the intermediary pools within a given loading path. Because large pools would most likely be found outside the vascular tissues – the latter being of relatively limited volume – it is assumed that their probable location is in the mesophyll. Our results regarding t0.5 values therefore suggest that at the lowest applied sucrose level IAA was being loaded primarily along a pathway passing through the protoplasts of mesophyll cells. At higher sucrose levels, the loading path was drastically diminished in size, suggesting that IAA loading was largely restricted to a direct transfer through the apoplast to the phloem, without passage through the mesophyll.

### Inhibition of phloem transport by sulfhydryl reagents

Photosynthates in leaves are generally loaded into the sieve element/companion cell complex through the plasma membrane from the apoplast or, alternatively, pass from the mesophyll to the phloem of minor veins through a symplastic pathway. Pathways may combine, run in parallel, or include a diffusive component depending on the species and on the physiological conditions within the tissue (Rennie and Turgeon, 2009). Evidence for the apoplastic loading of sugars and amino acids into the phloem has been provided for many plant species by testing their sensitivity to PCMBS, a membrane-impermeant inhibitor of proton-coupled transport (Lalonde et al., 2003). In *Ricinus* cotyledons externally applied [^14^C]sucrose was shown to move to the sieve elements in two parallel pathways, directly from the apoplast and indirectly after transit through the mesophyll cells (Orlich and Komor, 1992). One of the *Ricinus* sucrose carriers expressed in yeast can be inhibited by PCMBS (Weig and Komor, 1996).

The transport of simultaneously applied [^3^H]IAA and [^14^C]sucrose was measured with or without PCMBS or membrane-permeant p-chloromercuribenzoate (PCMB) to estimate the active, carrier-mediated component in their uptake. In the collected phloem exudate a steady concentration level for both [^3^H]IAA and [^14^C]sucrose was reached in all plants about 40 to 50 min after injection (Fig. 7). As judged by these equilibrium levels, the presence of PCMBS caused significant reductions in the active, carrier-based uptake of both [^3^H]IAA and [^14^C]sucrose by about 25% and 40% respectively (Fig. 7A and B). The observed responses suggest that in the loading pathway for IAA the active component is relatively smaller than that for sucrose. Alternatively, the two carriers may differ in their sensitivity to the inhibitor. However PCMBS also inhibits some aquaporins, which could upset water relations of the cells so altering the observed responses.

**Fig. 7.**
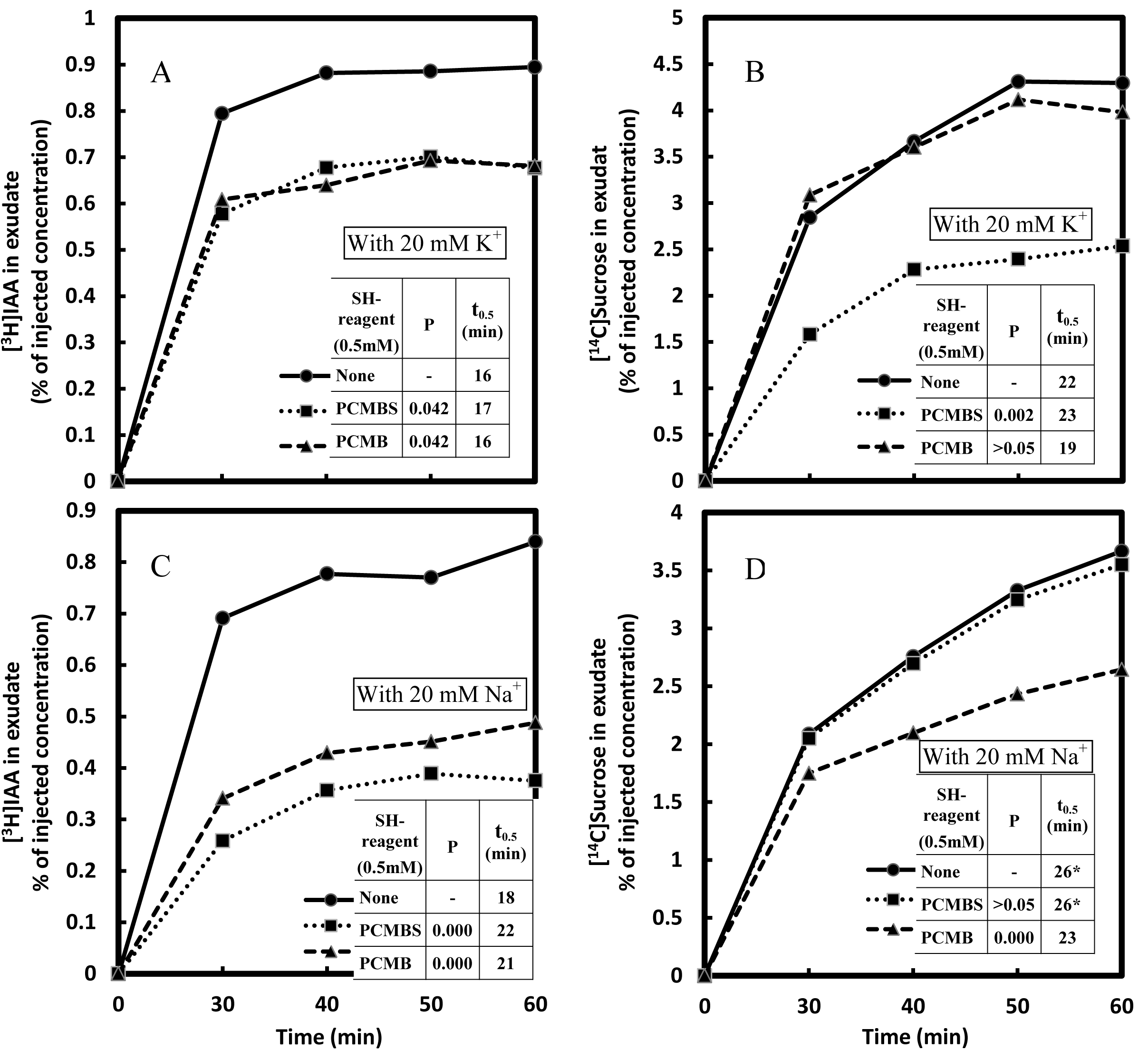
Effect of the sulfhydryl (SH) reagents PCMBS and PCMB on the transport of [^3^H]IAA (A and C) and [^14^C]sucrose (B and D). Each injection medium contained 30 mM MES buffer at pH 6.3, 0.73 MBq mL^−1^ [^3^H]IAA (0.79 μM), and 0.165 MBq mL^−1^[^14^C]sucrose (7.2 μM). The media also contained either 20 mM K^+^ (A and B) or 20 mM Na^+^ (C and D). Either 0.5 mM PCMBS or 0.5 mM PCMB was included in individual media, or no SH-reagent in the control. The data are combined from two experiments, and the means are from five replicate measurements. For other conditions or comments see Fig. 2.

With or without K^+^ present in the injection medium, PCMB inhibited IAA transport to, or nearly to, the same degree as did PCMBS (Figs. 7A, 7C; Na^+^ replaced K^+^ in 7C). In investigating uptake and movement of IAA in pea stems, Davies and Rubery (1978) found that whereas PCMBS decreased IAA accumulation in the stem segments, PCMB enhanced it. This was interpreted as penetrant PCMB blocking the IAA-efflux carrier on the interior side of the lower plasma membrane, so retaining more IAA in the transporting cells. That export into Ricinus phloem was inhibited by both PCMB and PCMBS is, however, not surprising even though carriers are clearly involved in IAA transport into the phloem: as the cut phloem where transport was measured involves an open ended system, any build-up in the transporting cells due to carrier disruption would simply remain in those cells and never reach the phloem. Nonetheless in the absence of K+ PCMB was slightly less effective an inhibitor than PCMBS, matching the promotion of phloem accumulation by TIBA.

When K^+^ was excluded from the injection medium (with 20 mM Na^+^ substituted for K^+^ in the buffer), the inhibitory effect of PCMBS on IAA entry into the phloem was about 54% (Fig. 7C), more than twice the effect obtained with K^+^ present (Fig. 7A). Therefore, the presence of 20 mM K^+^ was inhibitory for the active component in IAA loading. Interestingly, potassium ions had the opposite effect on sucrose loading: in the absence of K^+^, PCMBS was wholly ineffective against sucrose transport (Fig. 7D). Perhaps in the latter case the active component of sucrose uptake was being disabled by the low proton motive force caused by the sharply reduced availability of K^+^ for charge compensation (Malek and Baker, 1978). However the active loading of IAA not only continued, but actually doubled in rate when potassium ions were withheld from the injection medium. This could be explained if the processes of IAA and sucrose loading are driven by metabolic energy derived from two distinct sources.

While PCMBS was only effective in reducing sucrose transport into the phloem with K^+^, PCMB was only effective in the absence of K^+^ (Figs. 7B, 7D; Na^+^ replaced K^+^ in 7D). The efficiency of each of the inhibitors may be differentially affected by the prevailing proton motive force that is expected to vary with the applied K^+^ concentration (see above). The observed effects of K^+^ on sucrose loading may involve the regulatory activity of K^+^ channels located in phloem cells together with H^+^ pumps and sucrose carriers. The loss of AKT2/3 K^+^ channel function in an *Arabidopsis* mutant has been shown to result in impaired sucrose/H^+^ symporter activity and diminished phloem electric potential (Deeken et al., 2002).

### Effect of fusicoccin

The fungal toxin fusicoccin (FC) is widely used as a chemical probe in studies on active, energy-requiring processes. It can stimulate the activity of plasma membrane H^+^-pumping (P-type) ATPases in cells by binding to a 14-3-3 receptor protein (Sze et al., 1999), thereby affecting carrier-based proton-solute cotransport of various kinds, including sucrose loading in phloem. FC has been shown to enhance phloem transport of sucrose in *Ricinus* (Langhans et al., 2001; Malek and Baker, 1978) and in *Vicia* (Delrot and Bonnemain, 1981). In the experiment described here we investigated the effect of FC on phloem loading in *Ricinus* cotyledons of both sucrose and IAA simultaneously. The transport of [^14^C]sucrose, applied at 7.9 μM, was measured together with either 1.3 or 10.3 μM [^3^H]IAA, in the presence or absence of FC (0, 1, or 10 μM). At 1.3 μM IAA, sucrose transport was enhanced by FC (Fig. 8B), whereas in the same plants IAA transport was inhibited by FC (Fig. 8A). However, with 10.3 μM IAA present, FC failed to evoke these responses (Fig. 8C, D) as though IAA at the higher level was able to supersede or mimic FC’s action by evoking a parallel or identical effect. At 10.3 μM IAA, with no FC added, the relative rate of sucrose transport was doubled compared to the rate at 1.3 μM IAA (Fig. 8B, D). The enhancement was equal to that with 10 μM FC at the lower IAA level (Fig. 8B). Some of the responses to FC described here could have been affected by IAA in complex ways. The IAA-accelerated acidification of the apoplasm, at least in cells undergoing expansion, appears to be mediated by 14-3-3 receptors, transduction proteins not unlike the receptor for FC on the H^+^-ATPase enzyme (Sanders and Bethke, 2000; Trewavas, 2000). Alternatively, the amount of plasma membrane H^+^-ATPase may be increased by IAA (Hager et al., 1991).

**Fig. 8.**
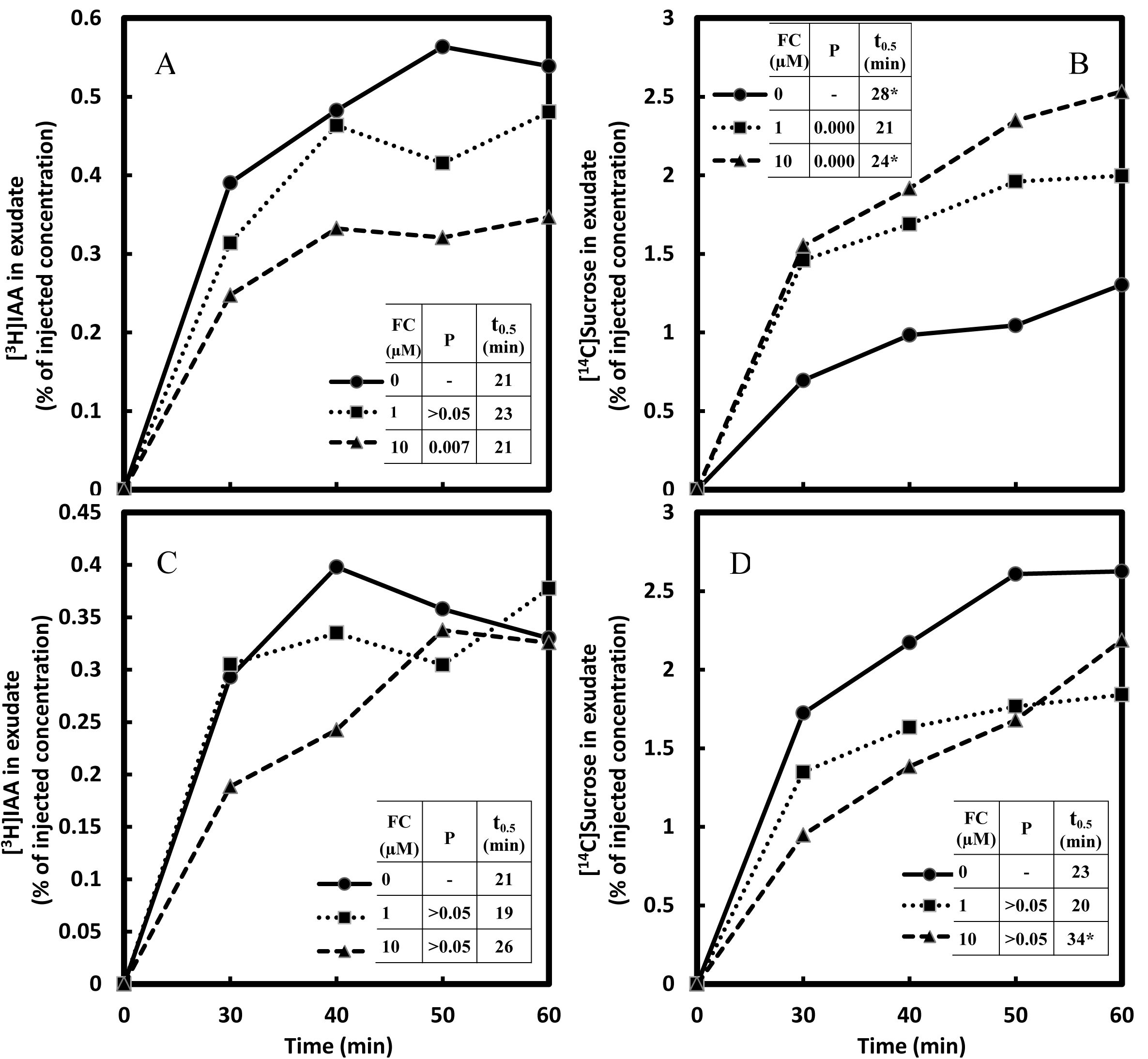
Effect of fusicoccin (FC) on the transport of [^3^H]IAA (A and C) and [^14^C]sucrose (B and D) at different IAA levels. Each injection medium contained 30 mM MES buffer at pH 6.3, 20 mM K^+^, 0.273 MBq mL^−1^ [^3^H]IAA, at either 1.3 μM IAA (A and B) or 10.3 μM IAA (C and D), and 0.18 MBq mL^−1^ [^14^C]sucrose (7.9 μM). FC concentrations were 0, 1, and 10 μM. The data are combined from two experiments, and the means are from five replicate measurements. For other conditions or comments see Fig. 2.

## Conclusion

In germinating *Ricinus* seedlings both sucrose and IAA derived from the endosperm are transferred into the peri-cotyledonary space and taken up by the cotyledons en route the seedling axis. This would involve uptake by the cells of the cotyledons and then cell-to-cell transfer to the companion cells of the phloem. Alternatively, movement at some point may be apoplastic prior to transfer into the companion cells. Sucrose has been reported to use both these routes. The synthetic auxin NAA competitively inhibited the IAA accumulation in the phloem showing that IAA was in part moving via auxin-specific transporters. PCMBS, which would act exterior to the cell membranes, reduced uptake of both IAA and sucrose by about 25% and 40%, respectively, indicating that carrier-mediated uptake into cells, not surprisingly, is involved at some point en route, and was more important for sucrose than for membrane-permeant IAA. As the IAA efflux-carrier inhibitor TIBA enhanced IAA accumulation in the phloem it would appear that the blocking of cell to cell IAA transport may force more IAA into the phloem, or that there is an efflux carrier sieve tubes themselves preventing diversion to other cells en route. The presence of K^+^ at low sucrose concentrations doubled IAA loading into the phloem, whereas at 100mM sucrose the loading of IAA was severely diminished in the presence of K+ even though sucrose without K+ had no effect. Thus the degree of sensitivity of IAA loading to the depolarization of the plasma membrane by K+ is correlated with sucrose concentration (the saturable influx via the proton cotransport system has a Km around 25 mM in Ricinus cotyledons though the value for the outer layer is about 5 mM (Komor, 1977)). At the lowest applied sucrose level, IAA was being loaded primarily along a pathway passing through the protoplasts of mesophyll cells, but at higher sucrose levels IAA loading appeared to be restricted to a direct transfer through the apoplast to the phloem, without passage through the mesophyll. We conclude that the transport of IAA into the phloem is multi-faceted with a carrier-mediated pathway playing a significant role.

## Materials and methods

### Plant material

Castor bean (*Ricinus communis* L. cv. Sanguineus) seeds, (Stokes Seeds, Inc. Buffalo, NY), were sown in a soil-less medium ‘Cornell Mix’ (Boodley et al., 1982), composed of peat, vermiculite and perlite at a ratio of 3:2:1, with supplements of fertilizers and pulverized dolomitic limestone. Seedlings were raised in a growth chamber under fluorescent lights with daily illumination for 16 h at 40 μMoles m^−2^ s^−1^. Day/night temperatures were 26/17 ^o^C. Six to seven-days-old seedlings with a well-developed endosperm were selected for treatment at a stage preceding the emergence of the cotyledons from within the endosperm. The seedlings were gently lifted from the growth medium so to minimize damage to the roots, were rinsed with deionized water, and hydrated for about two hours between layers of absorbent tissue paper that were moistened with 1 mM CaCl2 (Kallarackal et al., 1989)

### Injection media (IM)

#### Simultaneous transport of radiolabeled IAA and sucrose

IAA and sucrose transport were directly compared in individual seedlings using an injection medium in which defined levels of [^3^H]IAA and [^14^C]sucrose were combined in 30 mM MES buffer at pH 6.3. Either KOH or NaOH was used for pH adjustment in the MES stocks, giving a terminal concentration of 20 mM K^+^ or Na^+^ in the IM. Radioactivity levels were generally in the range of 0.5 to 1.5 MBq mL^−1^ for [^3^H]IAA, and 0.2 to 0.7 MBq mL^−1^ for [^14^C]sucrose, with corresponding concentrations averaging about 0.8 μM IAA and 8 μM sucrose. In some experiments, the concentration of IAA was adjusted to several different levels in a set of IM preparations by adding non-radioactive IAA, while keeping their [^3^H]IAA content the same. Comparable experiments were also done with sucrose. Actual radioactivity and concentration levels, together with notes on other components (e.g., specific chemical probes), if any, are provided with the individual figures. [^3^H]IAA (SA = 925 GBq mmol^−1^) was purchased from American Radiolabeled Chemicals (ARC) (Saint Louis, MO). [^14^C]sucrose was obtained from Sigma (Saint Louis, MO) (SA = 20.9 GBq mmol^−1^), or from ARC (SA = 22.9 GBq mmol^−1^).

### Incubation buffer (IB)

IB was used to incubate the endosperm during the transport experiment. It contained the same buffer as the one present in the corresponding IM injected *within* the endosperm, but not including any radiolabeled substances or chemical probes.

### Injection, transport, and recovery of radiolabeled substances

In preparing the seedlings for injection, the hypocotyl was cut with a sharp razor blade to remove the roots and lower hypocotyl, thus leaving a hypocotyl stump, seven to ten mm in length, attached to the cotyledons enclosed within the endosperm. Using a microsyringe, five μL IM was injected into the endosperm cavity between the cotyledons, thus exposing the two enclosed cotyledons to the radiolabeled substances being tested (Fig. 1). Then, the endosperm was placed in a small beaker, between layers of absorbent paper moistened with IB, ensuring that the endosperm with the emerging hypocotyl stump was held in an upright position. Freely exuding phloem sap from the cut surface of the hypocotyl was collected during ten minute intervals, starting at 0-10 minutes, generally for an hour, with a graduated microcapillary tube resting on the hypocotyl stump. High relative humidity was maintained throughout the transport period. The volume of the collected exudate was recorded, and the sample was transferred with 95% ethanol as a rinse into a liquid scintillation vial for analysis.

### Radioactivity Counting and Data Presentation

The ^3^H-and ^14^C-activity in each exudate sample was determined simultaneously using Ecoscint (National Diagnostics, Atlanta, GA, USA) with a Beckman (Fullerton, CA, USA) LS 1801 liquid scintillation counter programmed for dual-isotope DPM analysis. From the radioactivity of each isotope, the concentration of the respective transported substance was calculated. Specific activities used for the conversion were generally those provided by the manufacturer (with activities of [^3^H]IAA corrected for decay). Specific activity values were recalculated for those IM preparations in which a radiolabeled substance was supplemented with the corresponding non-radioactive compound. For the purpose of data evaluation and presentation, the concentration of a transported substance in each sample was expressed as a percentage of the concentration in the corresponding IM preparation; the data represent each individual 10 min collection and are not cumulative. The variable *t0.5* is equal to the time, in minutes, required for the transported substance to attain or approach one half of the steady state level in the phloem exudate, or one-half of the highest concentration obtained within a transport period of one hour in case a steady-state had not been fully attained. For plots not showing a tendency toward a steady-state, and/or remaining at or near linearity throughout the transport period, the *t0.5* value is marked with an asterisk (*) indicating a failure to attain a steady-state; the value of is then based on one-half of the highest level attained within 1 h of transport.

### Statistical Analysis

All of the data presented here are the combined results of at least two independent experiments, and represent the means of at least five replicate measurements. The number of replicates and of repeated experiments is stated in each figure caption. Data analysis was done with the Windows-based statistical system *Minitab* (Minitab, Inc.). To test the significance of each treatment effect compared to the relevant control, we calculated their *F* ratio along with the corresponding probability value using *Analysis of Covariance*, a model that allows for the correction of variation due to selected experimental conditions (e.g., sampling time, developmental variation, etc.) as appropriate. Significant treatment effects, thus corrected, were those within the probability range of P = 0.00 to 0.05.

## Acknowledgements

We thank the late Roger Spanswick and Robert Turgeon for constructive comments.

## Author contributions

IT developed the concepts, carried out the work and analyzed the data; IT and PD wrote the manuscript.

